# A rapid workflow for neuron counting in combined light sheet microscopy and magnetic resonance histology

**DOI:** 10.1101/2023.05.17.540884

**Authors:** Yuqi Tian, G Allan Johnson, Robert W Williams, Leonard White

**Affiliations:** Duke Center for In Vivo Microscopy, Department of Radiology, Duke University, Durham, NC, USA; Department of Genetics, Genomics and Informatics, University of Tennessee Health Science Center, Memphis, TN, USA; Department of Neurology, Duke University, Durham, NC, USA

**Keywords:** Neuron density, light sheet microscopy, mouse brain, neurologic image analysis, cell counting, neurons, three dimensional (3D)

## Abstract

Information on regional variation in cell numbers and densities in the CNS provides critical insight into structure, function, and the progression of CNS diseases. However, variability can be real or can be a consequence of methods that do not account for technical biases, including morphologic deformations, errors in the application of cell type labels and boundaries of regions, errors of counting rules and sampling sites. We address these issues of by introducing a workflow that consists of the following steps: 1. Magnetic resonance histology (MRH) to establish the size, shape, and regional morphology of the mouse brain in situ. 2. Light-sheet microscopy (LSM) to selectively label all neurons or other cells in the entire brain without sectioning artifacts. 3. Register LSM volumes to MRH volumes to correct for dissection errors and morphological deformations. 4. Implement novel protocol for automated sampling and counting of cells in 3D LSM volumes. This workflow can analyze the cells density of one brain region in less than 1 min and is highly replicable to cortical and subcortical gray matter regions and structures throughout the brain. We report deformation-corrected neuron (NeuN) counts and neuronal density in 13 representative regions in 5 C57B6/6J and 2 BXD strains. The data represent the variability among cases for the same brain region and across regions within case. Our data are consistent with previous studies. We demonstrate the application of our workflow to a mouse model of aging. This workflow improves the accuracy of neuron counting and the assessment of neuronal density on a region-by-region basis, with broad applications in how genetics, environment, and development across the lifespan impact brain structure.

## 1 Introduction

Accurate counts of neurons are essential metrics for understanding the structure and function of the brain (Bonthius et al. 2004; Herculano-Houzel and Lent 2005). This is particularly important for preclinical studies of human disease, as it provides a necessary starting point for assessing neurodegenerative changes on a regional basis(Bandeira, Lent, and Herculano-Houzel 2009) (Price et al. 2001) (Rosen and Williams 2001). In addition to absolute counts, measuring neuron density can provide important insights into the structural complexity of local circuits. The distribution of neurons in the mouse brain on a region-by-region basis is also crucial for understanding variation between individuals and strains. The density of neurons can reflect the distinctive cytoarchitecture of brain regions, including laminar organization, size and shape of constituent neurons, and the volume and composition of associated neuropil(Kasthuri et al. 2015) (Spocter et al. 2012) (Wree, Schleicher, and Zilles 1982). Furthermore, assessing neuronal densities in targeted brain regions under varying conditions may reveal impacts that change the composition of neuropil with or without associated loss or gain of neuronal cell bodies (Amunts et al. 1996; Selemon and Goldman-Rakic 1999). The heterogeneity of neuronal density within a given region can provide insights into the complexity, developmental history, and functional diversity of the region (Peters, Josephson, and Vincent 1991). Establishing such datasets enables pre-clinical studies of the impact of genetics, environment, and development across the lifespan on brain structure.

To investigate cells numbers and densities in CNS, researchers have developed various quantitative methods, including several stereological approaches (Williams and Rakic 1988) (West 1999; von Bartheld 2002) (Deniz et al. 2018) (Schmitz and Hof 2005) and approaches that first homogenize brain tissue and dissociate brain cells (Herculano-Houzel and Lent 2005) (Collins et al. 2010) (Young et al. 2012). Stereology is a quantitative method for estimating the three-dimensional characteristics of biological structures using two-dimensional histological sections or images and systematic random sampling within delineated brain regions. In traditional approaches to stereology, the specimen is preserved by means of chemical fixation, the brain is removed from skull, sectioned into thin slices (usually 5–50 μm thick), and stained with specific dyes to highlight different cell types. The sections are then viewed under a microscope, and measurements are taken to estimate the cell numbers within delineated brain regions. Both 3D counting and the optical fractionation are alternatives to traditional stereological methods introduced by Abercrombie (1946, reviewed in Williams and Rosen, 2003) that is design-based in its approach to sampling a volume of tissue, counting targeted objects (e.g., neurons), and generating statistics that assess the precision of the counts(West 1999). These methods considered unbiased by sectioning artifact, but still can suffer from bias due to differential z-axis shrinkage and variable penetration of stains. By providing a less biased estimate of structural parameters, these modern stereological methods have contributed significantly to our understanding of many biological systems. However, there are some limitations. Sterelogical methods require extensive preparation of tissue and the use of microtomes to section material. This can introduce damaging artifacts, lost sections, and distortions, even in the hands of experts. The results are sensitive to a range of factors, including the choice of chemical fixation, the post-fixation treatment of the specimens, the specific histological protocols used, and the specific features of the design-based approach to sampling and counting. Thus, it has been difficult to determine the accuracy of the measurements obtained in tissue that was subject to such deformation and the range of operational variables employed. Finally, stereological analysis can be a time-consuming process. For instance, the preparation and counting of brain tissue samples from the neocortex and hippocampus can take up to five days (Bonthius et al. 2004). When multiple brain regions are analyzed, the time and effort required can increase significantly, and the manual aspect of the process can introduce additional sources of variability.

In contrast, the isotropic fractionator (IF) involves homogenizing a small tissue sample from a region of interest—or the entire brain itself—and processing it into a uniform suspension of cells. The cells are then stained with a dye and placed into a special counting chamber, where the number of cells in a known volume is counted under a microscope(Herculano-Houzel and Lent 2005), or by the flow cytometry(Collins et al. 2010) (Young et al. 2012). The total number of cells in the brain region can then be estimated by extrapolating the cell density from the counting chamber to the reported or measured volume of the entire brain region. The downsides to isotropic fractionator are similar to stereology. One limitation is that both IF and stereology use the brain which is taken out of the skull and processed, introducing swelling or shrinkage, which can affect the accuracy of volume estimation;

Light sheet microscopy (LSM) has emerged as a powerful tool for 3D visualization of targeted populations of cells in intact, cleared tissue samples using fluorescent immunocytochemistry. Recent advances in tissue clearing (Ueda et al. 2020) have enabled the capture of high-quality, countable images in the whole-brain using LSM (Hillman et al. 2019). These intact 3D volumes can then be used for accurate counting of neurons and assessing neuronal densities, providing a more comprehensive and representative view of the brain region of interest. Typically, the whole brain images are registered to a standard reference atlas, such as the Waxholm Space Atlas (WHS) (Johnson et al. 2010) or the Allen Brain Atlas (ABA) (Wang, Ding et al. 2020). The combination of the digital images and the standard reference atlas facilitates the segmentation of regions and correction of altered brain morphology, allowing for comparison of neuron numbers and density within and across studies.

Researchers have employed this approach of registering LSM to WHS or ABA atlases in a number of recent studies (Renier et al. 2016) (Zhang et al. 2017) (Krupa et al. 2021) (Susaki et al. 2014) (Menegas et al. 2015). Susaki et al (Susaki et al. 2014) mapped LSM to WHS space but did not proceed with cytometric quantitative analysis. The remaining studies mapped the LSM images to ABA space for analysis. ClearMap, as described by Reiner et al (Renier et al. 2016), utilizes a peak detection algorithm with a threshold determined by comparing manual and machine counting, which may not yield optimal results when applied to complex and heterogeneous neuron distributions.

Zhang et al (Zhang et al. 2017) employed L1 minimization (Lasso), a machine learning regression method, to detect neurons in the whole brain. However, this method assumes sparsity of objects and a linear relationship between input features and output variables, with all features equally important, which may not hold true in the case of heterogeneous neuron distributions. Krupa et al (Krupa et al. 2021) used a 3D Unet for cell detection, but this method requires large inputs and considerable training time compared to machine learning. Menegas et al (Menegas et al. 2015) utilized Ilastics (https://www.ilastik.org) to segment cells in eight brain regions, with separate training for each region.

Registration of LSM images obtained from mouse brains to ABA space for correcting deformation and quantitatively segmenting brain structures and regions has one fundamental deficiency. The ABA atlas was generated from multiple specimens that were imaged after the brain was removed from the skull (Wang, Ding et al. 2020)). The lack of skull support and the effects of processing will deform the brain. Furthermore, ABA assembled their atlas from 1600 animals, which were registered into a volume that does not conform to the size and morphology of the *in situ* mouse brain—as we demonstrate in this work.

We present a novel approach that is highly reproducible and allows efficient counting across the whole brain. Our method utilizes magnetic resonance histology (MRH), an extension of MRI to microscopic resolution of fixed tissue specimens (Johnson et al. 1993). MRH images are acquired with the brain inside the skull, resulting in a representation that more closely approximates the size and morphology of the *in vivo* brain (Johnson GA 2023), as compared to reconstructions from histological methods that require dissection, serial sectioning of the brain, and chemical treatment of thin brain sections. As a result, MRH images can serve as the “gold standard” for correcting *ex situ* brain morphology. Our workflow offers a powerful combination of morphological correction based on MRH, machine learning-based neuron classification, and post-processing. It provides accurate and detailed neuron counts and calculations of neuronal density, which can be readily derived from different brain regions and specimens. Our method features a large, well-defined field of view in 3D, which allows for a comprehensive analysis of an entire subvolume under study. Moreover, our workflow applies the principles of optical fractionation to systematically sample and count neuron numbers in 3D volumes, which reduces the computational costs and represents a novel digital approach not previously explored.

## 2 Materials and Methods

### 2.1 Specimens

Two groups of animals were used to test the methods (see Table 1). The first group included 5 C57/B6 mice (4 male and 1 female) that were sacrificed at 90 ± 3 days to test the consistency of the counts and provide measures that could be related to existing literature. A second experiment with BXD89 mice included two male specimens at 111 and 687 days to test the sensitivity of the method to changes in neuronal density arising from aging. The BXD strains are a set of well-characterized recombinant inbred mouse strains, making them a valuable tool for systems genetics studies (Ashbrook et al. 2021).

**Table 1.** Test specimens for neuron counting.

| Specimen ID | Sex | Strain/Age |
| --- | --- | --- |
| 191209 | M | C57/90 d |
| 200302 | M | C57/90 d |
| 200316 | F | C57/90 d |
| 200826 | M | C57/92 d |
| 210823 | M | C57/93 d |
| 190108 | M | BXD89/111 d |
| 200803 | M | BXD89/687 d |

### 2.2 MRH, label map and LSM scanning

All animal procedures were carried out in accordance with guidelines approved by the Duke Institutional Animal Care and Use Committee. Specimens were perfusion-fixed using an active staining method as previously described (Johnson et al. 2019). Briefly, warm saline was perfused through a catheter in the left ventricle to exsanguinate, followed by ∼ 5 minutes of perfusion (50 ml) of a mixture of 10% buffered formalin and 10% Prohance (Gadoteridol) to reduce the spin lattice relaxation time (T1) of the tissue for accelerated scanning. MRH scanning was performed on a 9.4 T vertical bore magnet with a Resonance Research, Inc., gradient coil yielding peak gradients up to 2500 mT/m, controlled by an Agilent console running VnmrJ 4.0. The MRH scanning was performed using a Stesjkal Tanner spin echo sequence with b values of 3000 s/mm^2^ and 108 angular samples spaced uniformly on the unit sphere. Compressed sensing (Lustig 2007)was used with a compression factor of 8X (Wang et al. 2018), resulting in a large (252 GB) 4D volume with isotropic resolution of 15 um (Johnson GA 2023)

The label map in this study is based on a modified version of the Common Coordinate Frame (CCFv3) (Johnson et al. 2023) from ABA (Wang et al. 2020). The CCFv3 atlas includes 461 carefully curated regions of interest (ROIs). Our workflow relies on an initial mapping of labels from our canonical MRH atlas of a 90-day male C57/B6 mouse to the MRH of the specimen under study using a pipeline built around Advanced Normalization Tools (ANTs) (Anderson et al. 2019) (Avants 2011). Many of the CCFv3 ROIs are less than 1 mm^3^, including numerous subdivisions of cortical areas (laminae) and subcortical structures (subnuclei) that are of limited value within the present scope of our analyses. Accordingly, we reduced the label set to 360 ROI (180 per hemisphere) gray matter and white matter structures. This modified atlas, which we refer to as the reduced CCFv3 (r1CCFv3), consolidates these smaller subregions in CCFv3. The r1CCFv3 provides a label set that registers to the MRH volumes reproducibly enabling precise neuron counting within a large number of structures. The details of this registration process are described in (Johnson GA 2023; Johnson et al. 2022).

Following the MRH scans, the brains were carefully removed from the skulls, cleared using SHIELD (Park et al. 2019), stained using SWITCH (Murray et al. 2015), and scanned on a selective plane illumination microscope (SPIM) yielding whole brain images at a resolution of 1.8 × 1.8 × 4.0 um. The tissue clearing and SPIM imaging were performed at LifeCanvas Technology (Cambridge, MA). To transport the specimens to LifeCanvas, each specimen was immersed in neutral buffer with 0.5% Prohance in an airtight container. The container was then packed into a thermosafe box for transportation. The processing streams for MRH and LSM are depicted in Figure 1A.

**Figure 1.**
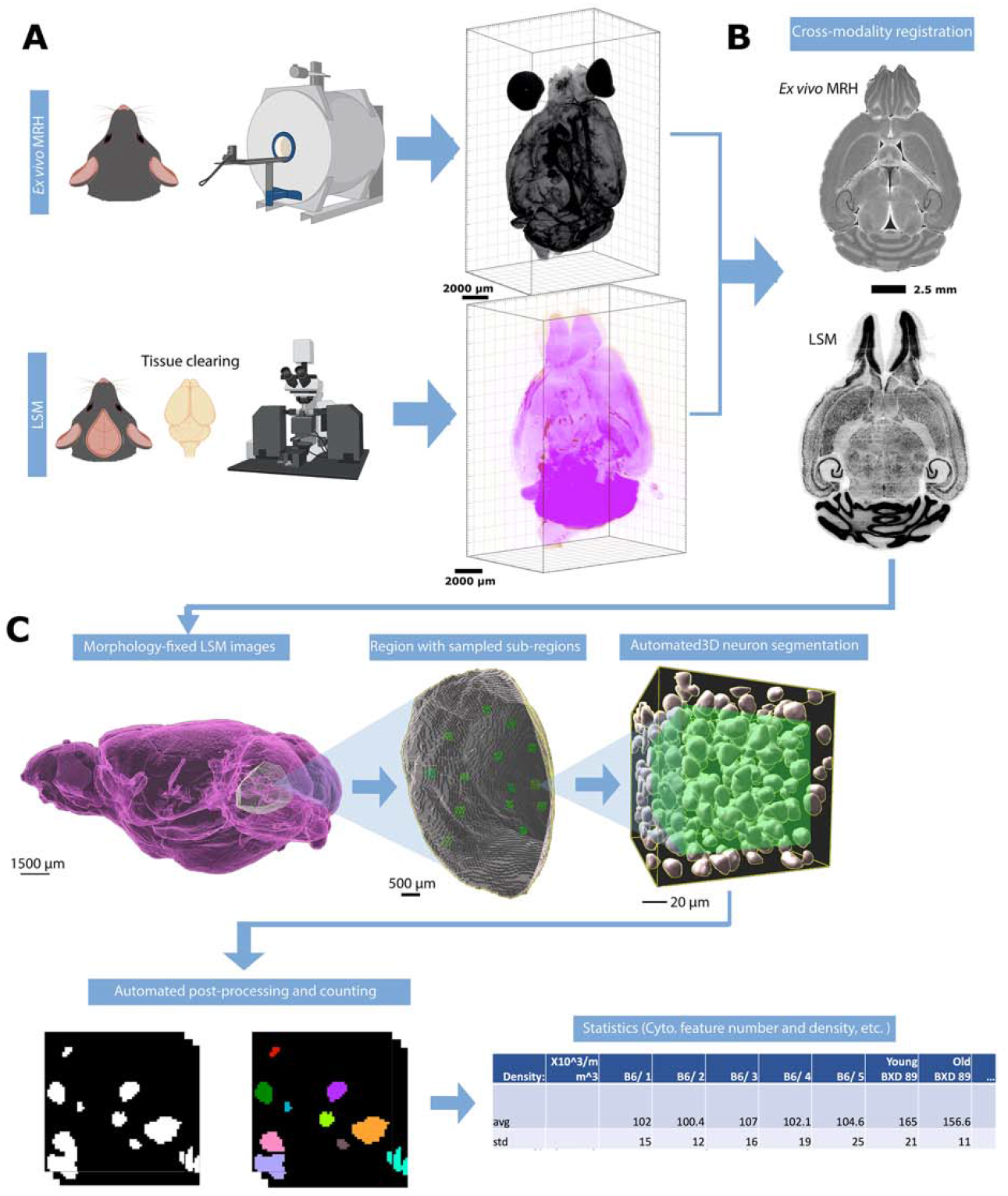
Overview of the workflow for assessing the density of neurons. (A) The mouse brain is imaged using two modalities: MRH imaging while the brain is in the skull, followed by LSM after the brain is removed from the skull and subjected to tissue clearing. (B) The LSM data are pre-processed by registering to MRH correcting the deformation in brain morphology. (C) The automated workflow locates the region with the label from r1CCFv3 and generates random subvolumes within that region to sample, applying the design-based principles of optical fractionation. Neurons in each subvolume are identified via a random forest algorithm followed by 3D watershed and volume filters and counted.

### 2.3 Automated Neuron counting

#### 2.3.1 Overview of the algorithm

To address the significant distortion introduced during skull removal and chemical processing in the preparation of LSM samples, the LSM images are first registered to the MRH volumes of the same specimen (Figure 1B) (Tian, Cook, and Johnson 2022). Section 3.1, accompanied by Figure 2, provides a detailed explanation of the impact of correcting the morphology and extraction of the cytometric statistics. We examine the effects of the preprocessing steps on the neuron density measurements, providing insights into the importance of these steps for accurate data analysis.

**Figure 2.**
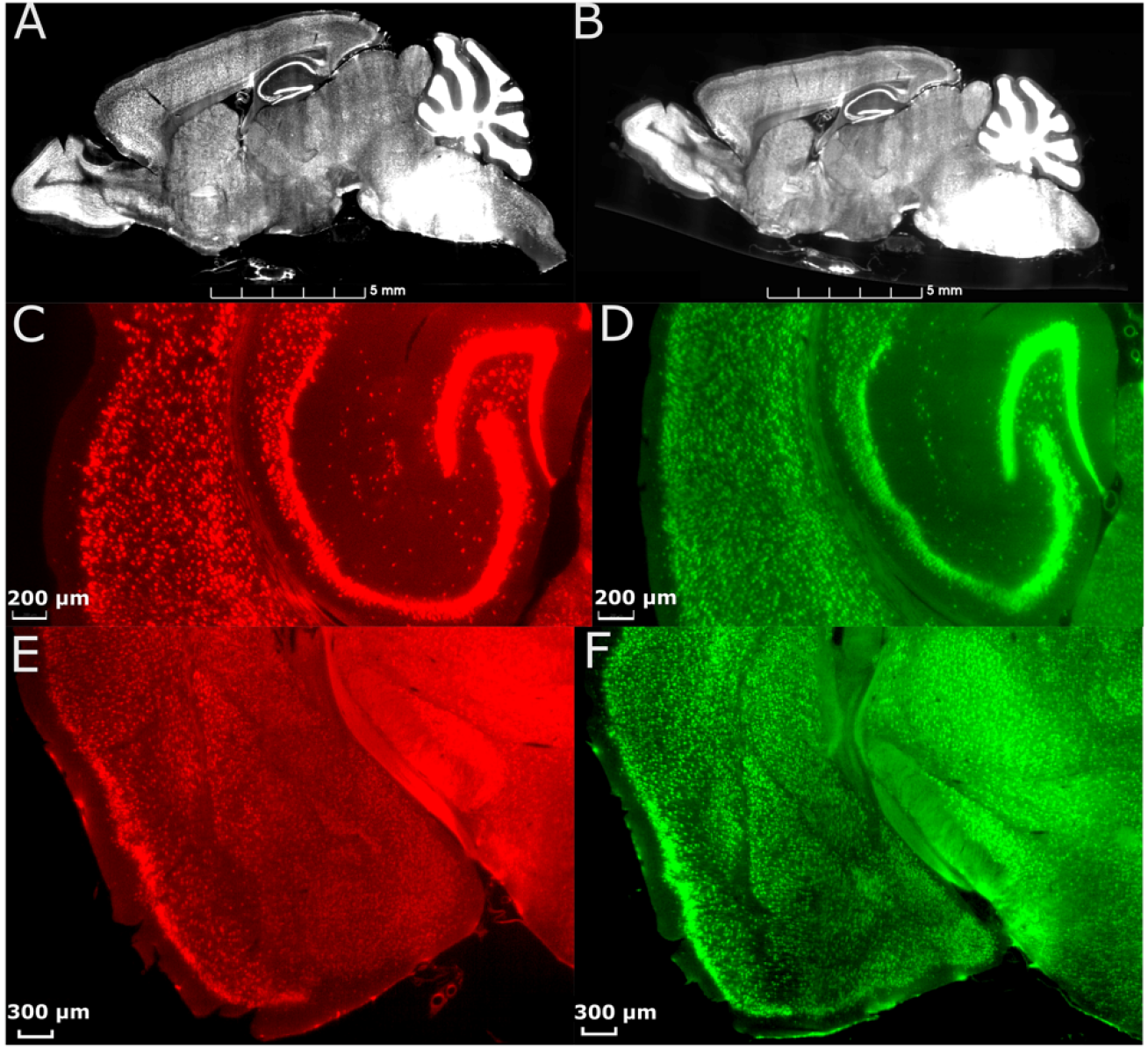
Effect of morphology correction on light-sheet datasets. (A) A single slice from a whole- brain LSM before correction. (B) The same slice as in (A) after correction using MRH. Magnified axial views of auditory areas and hippocampus before (C) and after (D) correction, and coronal views of the anterior temporal and posterior diencephalic region (showing the amygdaloid complex) before (E) and after (F) correction. The specimen is the same on both sides, and the colormap is used to illustrate the contrast between before and after correction.

The workflow is visualized in Figure 1C. After warping the LSM and label map to the MRH space, the user selects a region for analysis by inputting the corresponding label from the label map into Fiji. This generates a surface defining the region of the selected label, as well as a group of subvolumes within the surface. These subvolumes are distributed in a uniform but random manner, which improves the accuracy of neuron counting and facilitates analysis of the heterogeneity of neuron distribution throughout the selected region. For each subvolume, we apply a supervised algorithm called random forest neuron segmentation, which utilizes an ensemble of decision trees to generalize the projection between graphical features and labels. This algorithm is commonly used in image processing tasks and requires only a small amount of training data to perform well. Prior to segmentation, the random forest classifier is pre-trained using training data consisting of binary signatures, where neurons are labeled as 1 and non-neurons as 0, as well as feature vectors extracted from various visual characteristics of the image, such as texture, shape, and size. The training process optimizes the relationship between graphical features of an image and the assigned neuron labels.

The labeling example can be seen in Figure S1. After the random forest segmentation is applied to each subvolume, the workflow will perform post-processing on the objects classified as neurons. This post-processing uses 3D watershed and volume filters to further isolate the neurons as discrete objects. The final count of the neurons in each subvolume is generated based on these refined objects. The density is calculated by averaging subvolumes, and the standard deviation of the density provides insight into the variability of the neuron distribution within the region.

#### 2.3.2 Big data environment

Whole brain LSM images are large -- typically ∼ 300 GB. To enable efficient, interactive development and execution of the workflow we have assembled a hardware/software environment suited for these large data. The pipeline has been implemented on two high performance Dell servers: Dell E5-2400, NVIDIA Tesla V100 GPU and a Dell E52670, NVIDIA V100 GPU. Each server has 1.5TB of memory.

Several software packages have been integrated into the big data environment. Imaris (https://imaris.oxinst.com) is a commercial software package designed for interactive 3D/4D viewing of large microscopy data sets. It accommodates simultaneous visualization of multiple light sheet volumes using a hierarchical data format. Imaris has been developed to allow memory sharing with external packages. One of these is Fiji (https://fiji.sc/), a distribution of Image J that has been developed for applications such as that envisioned here through the extensive use of plugins. One crucial plugin for our work is BigDataViewer (Pietzsch et al. 2015), an open source solution for accommodating large volumes in Fiji and supporting plugins for post processing. This includes LabKit (Arzt M 2022) a user friendly Fiji plugin for microscopy segmentation using the random forest algorithm.

#### 2.3.3 Compiled algorithms

To improve the accuracy and efficiency of neuron counting, we developed two pipelines for different purposes: one for visualized validation and the other for production use. The initial pipeline (available on GitHub: [https://github.com/YuqiTianCIVM/NeuronCounting/blob/main/Algor_forVisualization.py]) provides visualization of classification and segmentation performance. This algorithm is built on the coding interface of the image rendering software, Imaris (https://imaris.oxinst.com). The procedures, from applying classifiers to obtaining statistics, are executed in the GUI of Imaris. Although the workflow cannot be fully automated through the Imaris coding interface, it can be automated through recording applications such as OS Automator (https://support.apple.com/guide/automator/intro-to-automator-aut6e8156d85/mac). The second algorithm (available on GitHub: [https://github.com/YuqiTianCIVM/NeuronCounting/blob/main/MainAlgor.py]) provides an automated workflow constructed using Python and macros. This method utilizes Fiji macro GUIs to automate segmentation, 3D watershed, and region counting plugins. The watershed uses morphological erosion to identify the center of each object, followed by calculating a distance map from the object center points to the object edges. The resulting topological map is then filled with imaginary water, and dams are built at locations where two watersheds meet to separate them. The region counting method assigns unique labels to each unconnected component to facilitate subsequent counting. The plugin used for constructing the random forest model is Labkit (https://imagej.net/plugins/labkit/). The watershed method and the 3D connected region counting method in the first algorithm are available in Imaris. The 3D watershed and 3D connected region counting method in the second algorithm are the Fiji/binary/watershed function and from the open-source plugin MorphoLibj (http://imagej.net/MorphoLibJ).

##### 2.3.3.1 Imaris pipeline for visualization

The user identifies the structure to be analyzed and loads the surface defining that structure. A Python script generates a collection of 3D sub regions that are randomly placed within the volume of the structure defined by the surface. Each counting sub region is a 100×100×100 μm cube. To avoid oversampling or undersampling in different regions and to accurately capture the heterogeneity, we typically analyze at least 15 subvolumes per brain region while keeping the number of subvolumes consistent across regions. The user selects “Labkit” in the Fuji extension. The GUI displays the sub-region’s volume, and the user either imports a pre-trained classifier or starts labeling neurons and background with binary labels. Figure S1 shows representative images of the training process. Once a classifier has been trained, it can be saved for future use in other ROI or other specimens, with some limitations discussed below. The classification is then sent to Imaris. Imaris applies thresholding to the binary labels and generates individual surfaces based on the results. The watershed algorithm uses seed points to split touching surfaces. After approximating the size of neurons through visualization, the volume filter is applied, and the count is generated.

##### 2.3.3.2 Fiji pipeline for high throughput

A Python script generates sub-regions inside the surface of the structure being analyzed and saves these subvolumes as individual TIFF files. The same Python script creates and executes a FIJI macro that reads the TIFF files in batches, applies the pre-trained random forest classifier, and performs 3D watershed. Components in the resulting image are labeled with different colors using the FIJI function “connected components labeling” and saved as a TIFF file. The second Python script reads these TIFF files in batches, applies volume filters, and generates counts of the components. The details of the volume filters are provided in Supplement Section 2.

### 2.4 Data availability statement

We have made the data in Table 1 available under a creative commons by NC-SA, accessible via this link: (https://civmimagespace.civm.duhs.duke.edu/login.php/client/4). The data is stored in H5 format, enabling interactive examination using Neuroglancer (https://github.com/google/neuroglancer). Readers can sign up to access the data, while reviewers may request additional anonymous accounts for access.

## 3 Results

### 3.1 Visualization of morphology correction

Figure 2 displays the comparison between images from an uncorrected and deformation-corrected whole brain light sheet volume. Much of the existing literature for stereology has employed optical light microscopy and confocal microscopy with higher resolution and smaller field of view than that in the whole brain LSM images. However, the deformation, which is dependent on the fixation and staining method, is likely to be similar to that presented in Figure 2A, C, E. The compensation of this deformation is usually done through the application of an isotropic correction factor, but this factor may be insufficient to fully address the nonuniform deformation arising from tissue processing.

Figure 2B shows the LSM image with morphology correction by MRH, which significantly reduces the irregular distortion present in the raw LSM images. The magnified views of the posterior cerebrum (showing the auditory cortex and hippocampus (Figure 2D) and LGN (Figure 2F) illustrate the benefits of the morphology correction method in improving the quality of imaging data and segmentation label maps, compared to Figure 2C and Figure 2E. Figure S2 illustrates the differences in neuron density between raw LSM and deformation corrected LSM for 6 brain regions. The smallest percent difference (calculated as the difference between raw and corrected density over the corrected density), observed in the primary visual cortex, is 7%, while the largest, observed in the subiculum, is 64%. These results demonstrate that deformation is not uniform throughout the brain.

Figure 3 A,C,E displays the label-map CCFv3. Data for the ABA were obtained by averaging 2-photon microscopy images of 1,600 animals. The distortions from tissue handling are not consistent across all 1600 animals. Wang et all report variability in the volume of the regions in their atlas ranging from 6 to 80% so tissue handling introduces variation (Wang et al. 2020). Figure 3 B,D,F displays the r1CCFv3 labels with morphology correction by MRH. The resulting corrected label map provides more accurate segmentation of brain regions.

**Figure 3.**
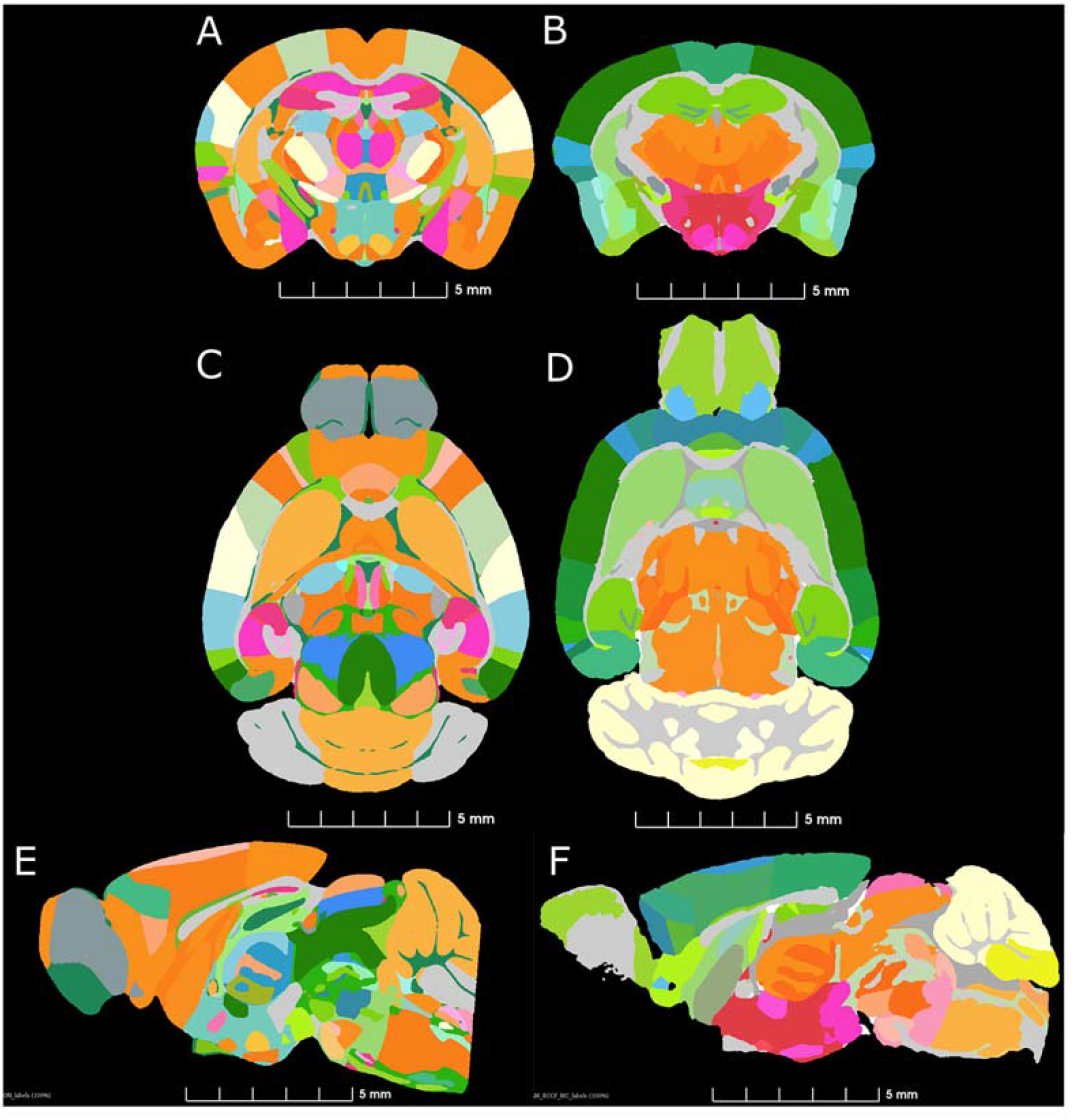
Comparison of similar levels from the ABA (A,C,E) and a single specimen imaged by MRH. (A,D,F). The colormaps are different because the ABA atlas is shown with the full complement of 461 (CCFv3) labels and the MRH is shown with the reduced (r1CCFv3) labels in which some ROI have been combined.

### 3.2 Accuracy of the / Comparison of machine counting and human counting

To assess the accuracy and precision of our workflow, we chose five brain regions with cell densities we determined would be countable: Dorsal part of the lateral geniculate complex (LGd), Auditory area (AUD), retrosplenial area (RSP), orbital area (Orb), and subicular region (SUBR). We created 15-20 3D sub volumes for sampling within these regions with dimensions of 1000 μm x 1000 μm x 12 μm (556 × 556 × 3 voxels) for visualization in Figure 4A and B, and 100 μm x 100 μm x 100 μm (56 × 56 × 25 voxels) for routine implementation of the workflow. The random forest classifier, trained on a random sub volume in the auditory cortex of a specimen not involved in the experiment, was applied automatically to identify the objects to be included. We randomly selected 10-15 sub volumes within the same datasets, and an experienced researcher manually counted the neurons within them by labeling neurons in sub volumes across 3-5 specimens. We compared the results of both methods and found that both manual counting and the machine had overall performance that was comparable (Figure 4A, B). A statistical comparison is shown in Figure 4C.

**Figure 4.**
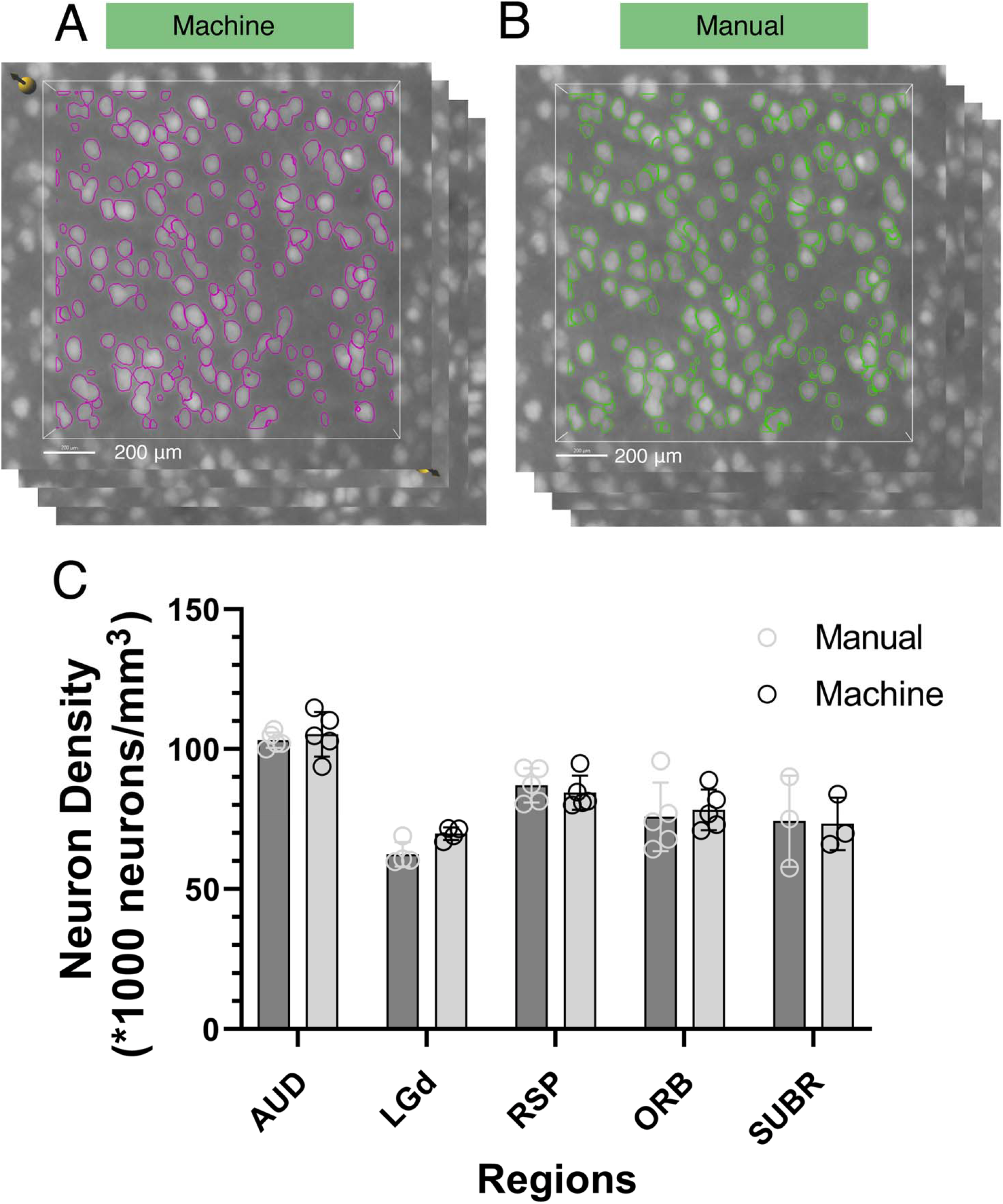
Comparison of neuron density between manual counting and the machine workflow. Standard deviation is shown, representing statistical variations across five different 90-day C57/B6 specimens.

### 3.3 Neuron counting in C57 mouse brain

We demonstrated the use of the pipeline in counting neurons in 13 different brain regions across 7 specimens. We choose representative regions to demonstrate the capability of our workflow in neocortex, hippocampus, amygdala, thalamus, and brainstem. These regions include (1) neocortex: orbital area (Orb), auditory area (AUD), retrosplenial area (RSP), primary visual area (VISp); (2) hippocampal region: entorhinal area (ENT), subicular region (SUBR), field CA1 (CA1), field CA3 (CA3); (3) amygdala: basolateral amygdalar nucleus (BLA); (4) thalamus: dorsal part of the lateral geniculate complex (LGd), and the entire thalamus (Th); (5) brainstem: facial motor nucleus (VII), principal sensory nucleus of the trigeminal (PSV). Initial measurements were performed on 5, 90-day old male C57/B6 specimens and revealed that the neuron density in some regions (LGd,AUD,RSP,Orb) remains quite consistent across specimens, while there is a high standard deviation in others (PSV,VISp,TH,BLA) (Figure 5). Figure S3 and Table S1 present the comparison between the neuron density obtained from our workflow and previous studies.

**Figure 5.**
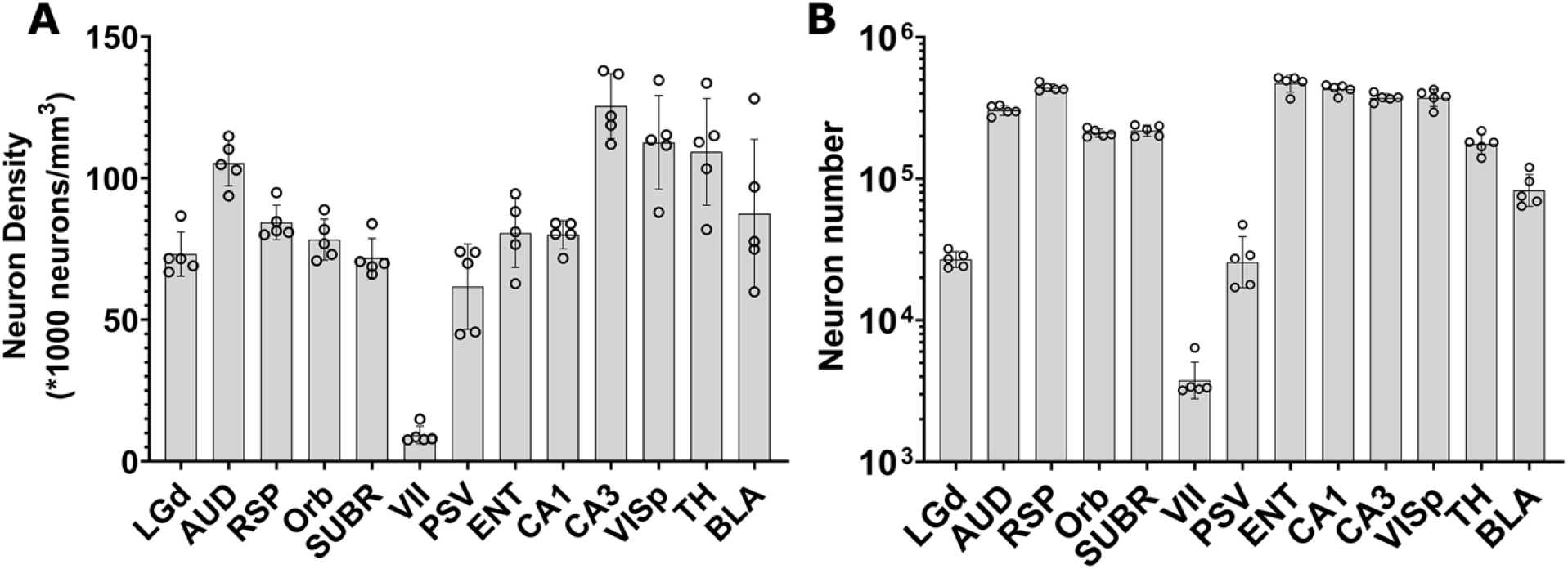
demonstrates the variation of neuron density (A) and neuron number (B) across brain regions, as well as variations across specimens. Error bars are standard deviations of the means. The dots indicate neuron counts from individual specimens. Region: LGd: Dorsal lateral geniculate nucleus, AUD: auditory area, RSP: retrosplenial area, Orb: orbital area, SUBR: subiculum, VII: facial motor cortex, PSV: trigeminal, ENT: entorhinal area, CA1: field CA1, CA3: field CA3, VISp: primary cortex, TH: thalamus, BLA: Basolateral amygdala

To assess the variability of our measurements, we calculated the coefficient of variation (CV) for the density and number of neurons in several brain regions across different specimens (Table S2). The CV for neuron density ranged from 0.062 in CA1 to 0.333 in VII, with an average of 0.139. The CV for neuron number ranged from 0.06 in CA1 to 0.44 in PSV, with an average of 0.17. These results indicate that there is considerable variation in neuron density and number across individual animals, even within the same brain region. Notably, the regions with the highest CV for neuron density and number were VII (0.333) and PSV (0.44), respectively. This is likely to be induced by the small sizes of these regions. The higher coefficient of variation for small regions may be due to minor displacements between the placement of individual delineations from the labelmap and the actual neuroanatomical structures, as discerned in LSM, especially for brainstem regions where deformation was significant. Conversely, the regions with the lowest CV for neuron density and number were CA1 (0.062), RSP (0.073), AUD (0.076), Orb (0.092), CA3 (0.091), and SUBR (0.097), respectively.

These findings underscore the importance of accounting for inter-individual variability when analyzing neuronal parameters and highlight the need for careful consideration of sample size and statistical power in studies of brain structure.

### 3.4 Illustration of aging effects on the neuron density and number

Figure 6 presents a comparison between young (111 day) and old (687 day) BXD89 specimens to examine the impact of aging on neuron density. It is important to note that the standard deviation in Figure 6 reflects the heterogeneity of neuron distribution within each brain region, which was obtained by analyzing sub volumes. Our analysis revealed that in certain regions, such as the orbital area (Orb), the observed decrease in neuron density was primarily due to a reduction in the total number of neurons, while in other regions such as the entorhinal area (ENT) and basolateral amygdala (BLA), the observed decrease in density was largely attributable to regional enlargement of the brain during aging. These findings underscore regional variation and the complexity of the aging process and highlight the need for careful consideration of volume and cellular number when studying brain development across the lifespan.

**Figure 6.**
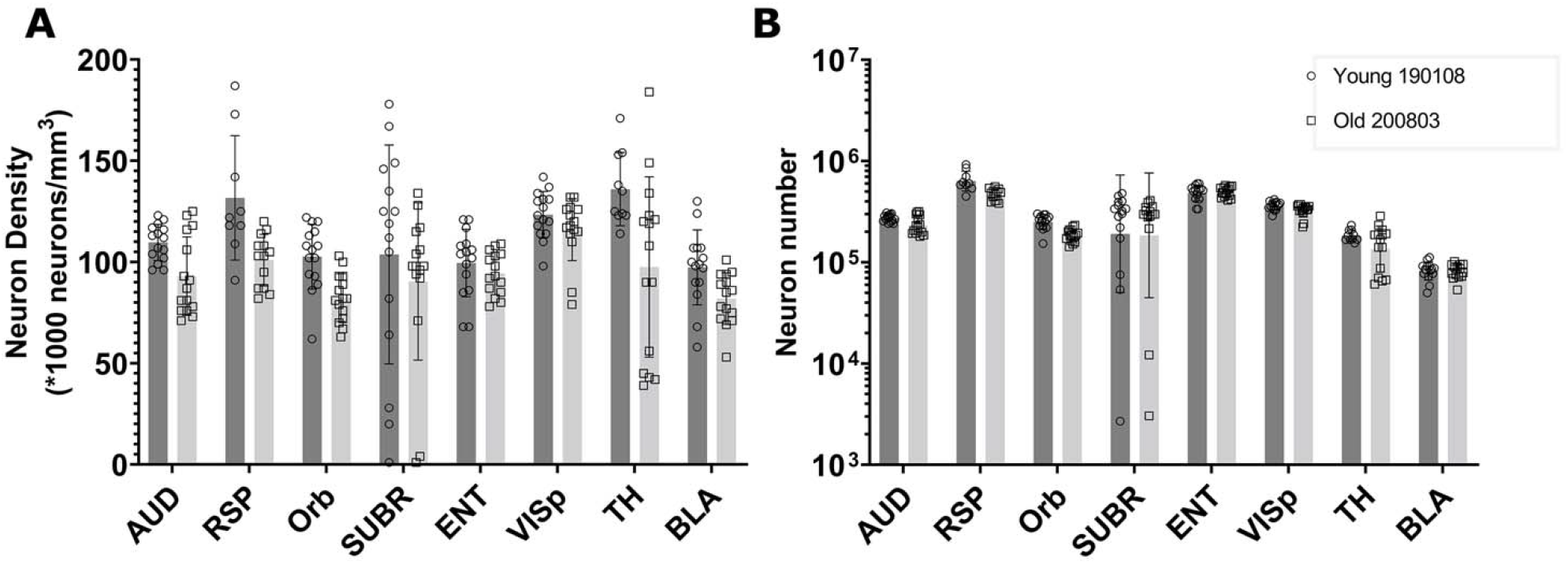
compares the neuron density and number in multiple brain regions of young and old BXD89 mice, highlighting the impact of aging on neuronal populations. The error bars in this figure represent the standard deviation of neuron distribution within each region, obtained from sub volumes.

## 4 Discussion

We present a workflow that corrects the deformation introduced by the chemical treatment of brain tissue for LSM and uses an automated algorithm for neuron segmentation and counting through machine learning and post-processing. To improve the accuracy of the results, we use the MRH of the same specimen as a reference for LSM image morphology, as MRH provides in-skull brain images that closely approximate in vivo brain images. Our method offers a reliable means of acquiring neuron number and density and requires less human labor and subjective judgment than previous methods. It is also much easier to replicate across multiple brain regions and specimens. By correcting raw images and conducting region segmentation, our method provides improved estimates over previous counting methods. Furthermore, by incorporating the principles of optical fractionation, our workflow provides an unbiased sampling method with equal probability for each section, resulting in reduced computational costs and representing a novel digital approach that has not been explored before.

To validate the method, we used the workflow to analyze 13 regions in different divisions of 5, 90-day C57/B6 male specimens. The results, as shown in Figure 5, demonstrate the variability in neuron density across regions within the same animal and across animals for the same regions. In the supplemental materials, we conducted a systematic search of the literature providing neuron counts of the same regions for comparison with our method. As our method produces regional neuron density as its direct output, we assessed the accuracy of our results in terms of density. For most cortical regions, our findings were consistent with those of previous studies or fell within the range of previously reported results. However, in some regions, e.g., LGd, PSV, VISP, TH, and BLA, our results showed a higher neuron density than what was reported in earlier studies. This is likely due to our utilization of the morphology-corrected space for calculating neuron density. This space closely approximates the volume and morphology of the brain confined within the skull, which may have contributed to the observed differences in density measurements. In certain regions where our density measurements were markedly different from those reported in previous studies, such as the facial motor nucleus, we manually counted the sub-regions to validate the neuron distribution. Our findings indicated that the neuron distribution in this region was indeed sparse, with a density measurement about six times lower than that reported in the Blue Brain Map (BBM) (Ero et al. 2018). We hypothesize that this discrepancy may be due to differences in data quality or acquisition methods between studies. BBM estimated neuron counts using a transfer function and Monte Carlo method applied to the ABA Nissl atlas. The authors only used whole-brain values from literature, such as the total number of cells and neurons in the mouse brain, to constrain their estimates of cell densities in each brain region. Further investigation is required to elucidate the causes of the observed discrepancy. Nonetheless, our method, which directly measures regional neuron density, provides a reliable alternative to indirect methods such as those used by BBM, and can contribute to a better understanding of brain structure.

Our method has two limitations. In those regions in which the cell density is so high that the resolution of the LSM is not sufficient to differentiate countable cells (e.g., dentate gyrus), the random forest classifier and watershed algorithms will fail. A second limitation is that performance is dependent on the quality of the raw data. In cases where the tissue preparation and scanning are suboptimal, the method will not function properly. Although the data provided by LifeCanvas, which we used for our study, is generally of high quality, there are occasionally regions with poor image quality (as shown in Figure S4), which can limit the method’s reliability. Advanced imaging techniques such as expansion microscopy, as demonstrated by ((Wassie, Zhao, and Boyden 2019) or acquisition with higher power objectives will address resolution questions and are likely to produce reliable and accurate counts of neurons using our workflow.

The application of stereology to counting cells has a venerable history. As acquisition methods have improved the methods to count cells have been adapted to the newer data. The advent of tissue clearing and volume light sheet imaging yield opportunities for significant improvement in precision and accuracy. This is particularly important in our studies of the mouse brain as we seek quantitative understanding in neurodegeneration and aging. Correction for tissue changes, use of a more accurate label set, and automation described in this work will provide crucial tools in quantitative neuropathology.

## 6 Author Contributions

YT generated the code performed the evaluation and wrote the manuscript. GJ conceived the project and assisted writing the manuscript. RW provided insight into stereology. LW provided insight into the importance of measuring neuron number and density. YT, GJ and LW formulated the strategy. GJ and RW provided funding and assisted in writing the manuscript. All authors contributed to the manuscript revision, read, and approved the submission.

## 8 Acknowledgments

We are grateful to James J. Cook for technical support and assistance in software development. We are grateful to Tatiana Johnson for assistance in editing the manuscript.

