## Supplementary Materials for "A rapid workflow for neuron counting in combined light sheet microscopy and magnetic resonance histology"

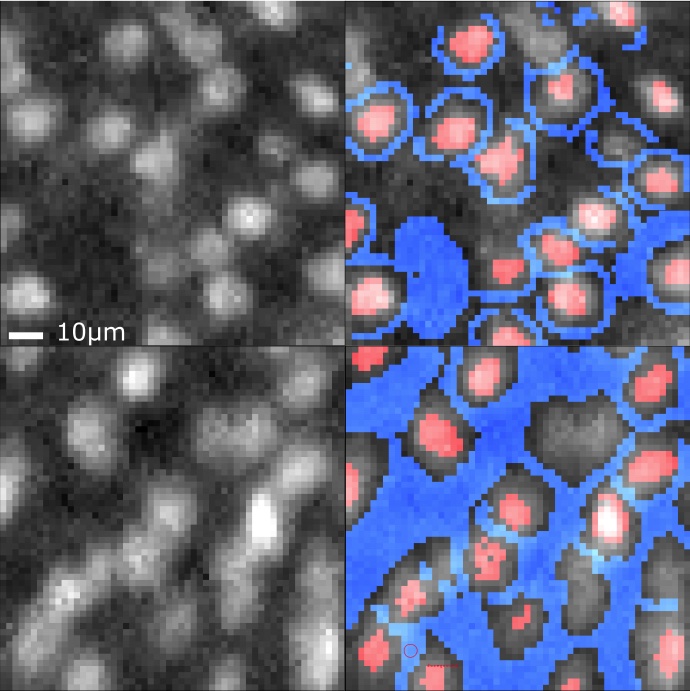

**Supplementary Figure 1.** Left: Raw light sheet images from one of the sub volumes used in the counting algorithm. Right: Training data with blue sampling background and red labeled cells. Typically, around 20 neurons need to be labeled with both the cell body and background for the algorithm to learn the accurate features of neurons. The figure depicts two different scenarios. In the first row (auditory cortex), there are no overlapping neurons, and the background labels are applied between the neurons and the empty spaces. In the second row (field CA3), the image shows connected neurons, and it's recommended to avoid adding a background label in the connected regions as it can lead to inaccurate classification. Instead, leave the connected neuron segmentation to watershed and volume filter.

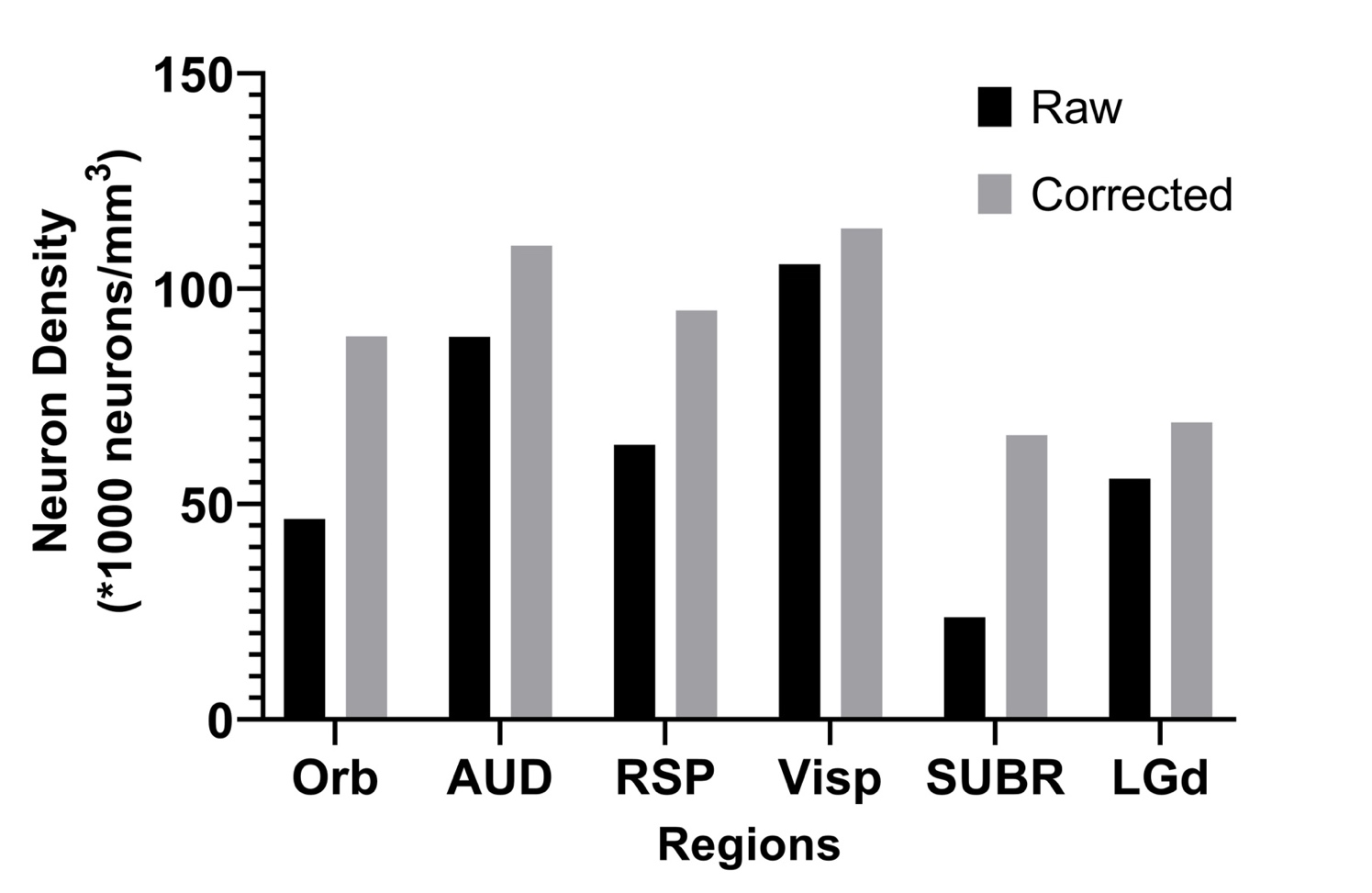

**Supplementary Figure 2.** shows the comparison of neuron density between raw LSM data and deformation corrected LSM data for six brain regions. Specimen: 200316.

**Table S1 Comparison with other works**

Figure S3 and Table S1 present a comparison between the neuron density measurements obtained from our workflow and those reported in other studies. As our workflow reports neuron density, we converted other studies' reported neuron numbers (if any) to neuron density by dividing them with the corresponding CCFv3 volume. Our results are generally consistent with those of other studies, but in some regions, such as LGd, TH, BLA, we observe higher neuron density. This difference may be attributed to the fact that we removed tissue swelling during LSM data pre-processing. In regions with high heterogeneity in neuron distribution, such as subiculum and ENT, different studies report varying neuron density values, and our results fall within that range. This suggests that our results cover sub-regions with both sparse and dense neuron distribution in these regions. The findings are illustrated in Figure S1.

| Region | Reference | (Kuronen, Lehesjoki et al. 2012) | (Dursun, Jakubowska-Dogru et al. 2011) | (Scott, Jeffrey et al. 1994) | (Seecharan, Kulkarni et al. 2003) | (Ero, Gewaltig et al. 2018) | This work |
| --- | --- | --- | --- | --- | --- | --- | --- |
| Dorsal lateral geniculate nucleus | **Neuron density(no/**$\boldsymbol{mm}^{\boldsymbol{3}}$**)** | 55965 | 44517 | 38158 | 66333 | 62465 | 73200 |
|  | **Reference** | (Trujillo-Estrada, Davila et al. 2014) | (Fabricius, Wortwein et al. 2008) | (Ero, Gewaltig et al. 2018) |  |  |  |
| Subiculum | **Neuron density(no/**$\boldsymbol{mm}^{\boldsymbol{3}}$**)** | 155500 | 46965 | 83717 |  |  | 71860 |
|  | **Reference** | (Herculano-Houzel, Watson et al. 2013) | (Ero, Gewaltig et al. 2018) |  |  |  |  |
| Entorhinal areas | **Reference** | 66988 | 91110 |  |  |  | 80640 |
|  | **Neuron density(no/**$\boldsymbol{mm}^{\boldsymbol{3}}$**)** | (Herculano-Houzel, Watson et al. 2013) | (Ero, Gewaltig et al. 2018) |  |  |  |  |
| Auditory | **Neuron density(no/**$\boldsymbol{mm}^{\boldsymbol{3}}$**)** | 109,730 | 108440 |  |  |  | 105805 |
|  | **Reference** | (Herculano-Houzel, Watson et al. 2013) | (Ero, Gewaltig et al. 2018) |  |  |  |  |
| Retroplenial | **Neuron density(no/**$\boldsymbol{mm}^{\boldsymbol{3}}$**)** | 98148 | 101650 |  |  |  | 84479 |
|  | **Reference** | (Herculano-Houzel, Watson et al. 2013) | (Ero, Gewaltig et al. 2018) |  |  |  |  |
| Orbital | **Neuron density(no/**$\boldsymbol{mm}^{\boldsymbol{3}}$**)** | 48109 | 77961 |  |  |  | 78333 |
|  | **Reference** |  | (Ero, Gewaltig et al. 2018) |  |  |  |  |
| Facial motor nucleus | **Neuron density(no/**$\boldsymbol{mm}^{\boldsymbol{3}}$**)** |  | 36427 |  |  |  | 9700 |
|  | **Reference** |  | (Ero, Gewaltig et al. 2018) |  |  |  |  |
| trigeminal | **Neuron density(no/**$\boldsymbol{mm}^{\boldsymbol{3}}$**)** |  | 45986 |  |  |  | 58607 |
|  | **Reference** | (Fabricius, Wortwein et al. 2008) | (Ero, Gewaltig et al. 2018) | (Hlatky, Lui et al. 2003) | (Liu, Yu et al. 2006) |  |  |
| CA1 | **Neuron density(no/**$\boldsymbol{mm}^{\boldsymbol{3}}$**)** | 44500 | 96289 | 52000 | 38290 |  | 80293 |
|  | **Reference** | (Fabricius, Wortwein et al. 2008) | (Ero, Gewaltig et al. 2018) | (Hlatky, Lui et al. 2003) |  |  |  |
| CA3 | **Neuron density(no/**$\boldsymbol{mm}^{\boldsymbol{3}}$**)** | 53900 | 135494 | 88000 |  |  | 112233 |
|  | **Reference** |  | (Ero, Gewaltig et al. 2018) |  |  |  |  |
| Primary visual | **Neuron density(no/**$\boldsymbol{mm}^{\boldsymbol{3}}$**)** |  | 100245 |  |  |  | 115343 |
|  | **Reference** |  | (Ero, Gewaltig et al. 2018) |  |  |  |  |
| Thalamus | **Neuron density(no/**$\boldsymbol{mm}^{\boldsymbol{3}}$**)** |  | 78611 |  |  |  | 108200 |
|  | **Reference** |  | (Ero, Gewaltig et al. 2018) | (Mozhui, Hamre et al. 2007) |  |  |  |
| BLA | **Neuron density(no/**$\boldsymbol{mm}^{\boldsymbol{3}}$**)** |  | 77920 | 74375 |  |  | 87386 |

**Supplementary Table 1.** The comparison of neuron density across literature and from our reported workflow.

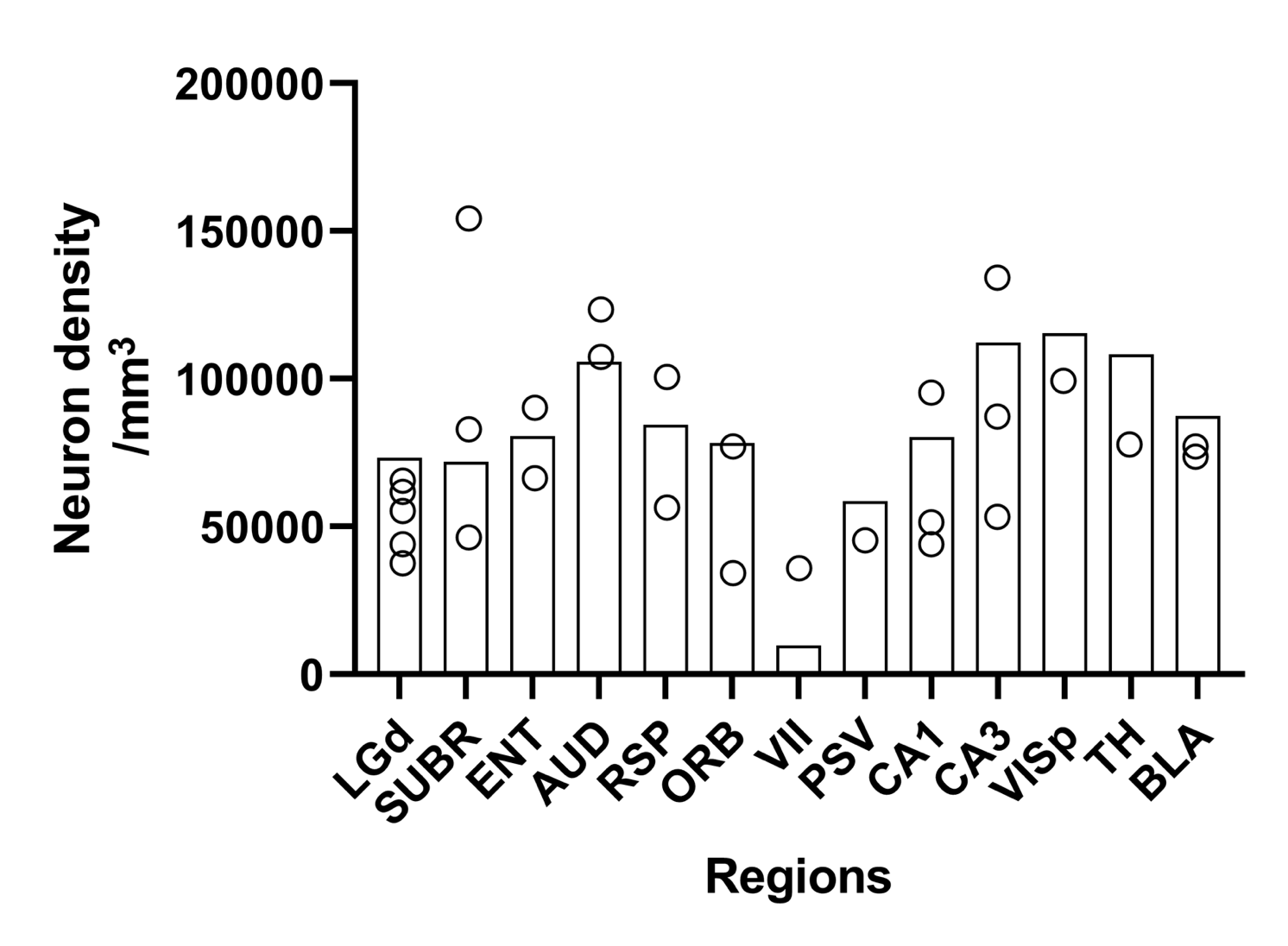

**Supplementary Figure 3.** presents a comparison of neuron density values obtained from our workflow with those reported in the literature. The figure highlights differences between our findings and those of other studies in different brain regions. The bar represents the values obtained from our workflow, while the circles indicate the corresponding values reported in peer studies.

**S1.1 Calculation of neural density across literature**

The literature search was conducted using keywords related to brain regions and their broader categories. The databases used were Google Scholar and PubMed. For example, in Entorhinal cortex, the keyword is below:

neuron* AND (densit* OR population* OR number* OR cell atlas) AND ("entorhinal cortex" OR EC OR allocortex) AND ("mice" OR "mouse")

During the literature search, if a study uses methods such as IF or stereology, and reports both the neuron number and the neuron density, we will divide the reported number by the corresponding CCFv3 volume.

Regarding subject strains, age, and sex, these factors can have an impact on neuron density. If a study presents the neuron density or number indirectly, we will provide further explanation. However, if a study does not report age or sex, no mention will be made in the explanation.

**LGd:** In the first three references, the neuron numbers are approximated from the figures they provided for the wild type (WT, C57/B6).

In (Kuronen, Lehesjoki et al. 2012), the number is the average of WT 1 month and 5 month.

In (Dursun, Jakubowska-Dogru et al. 2011), the number is from newborn control group.

In (Scott, Jeffrey et al. 1994), the number is from 100 days control group.

In (Seecharan, Kulkarni et al. 2003), age is not specified for 8 C57/B6 mice, but for the large herd of mice, the average age is 100d and sex ratio is 1:1. The neuron number is obtained from Table1.

As explained in main text, (Ero, Gewaltig et al. 2018) does not include the measurement but instead employs the whole brain cell counts and monte carlo method to approximate the distribution. The blue brain map from (Ero, Gewaltig et al. 2018) provides the density.

**Subiculum:** In (Trujillo-Estrada, Davila et al. 2014):

The neuron density is obtained from the equation: Neuron density ~= density of SoM-interneurons + PV-interneurons (inhibitory neurons) + Principal neurons (excitatory neurons). Therefore, the neuron density is approximated from the figures:

2 month WT (C57BL6) ~= 17000 + 10500 + 128000 ~ 155500 neurons/mm^3

6 month WT (C57BL6) ~= 12000 + 10500 + 120000 ~ 142500 neurons/mm^3

In (Fabricius, Wortwein et al. 2008): The neuron number is obtained from optical fractioner, and therefore we have to use the subvolume they provide to get the density. Control group (C57 2month 4 male and 8 female) mean neuron number is 181. Volume is obtained from Cavelieri principles. Volume = 2.51mm^2 x section thickness = 2.51mm^2 x 39um. The density is 46965/mm^3.

In (Ero, Gewaltig et al. 2018): They approximated the neural density from the observed density by a transfer function. The neuron density of subiculum provided in supplemental is 83717/mm^3.

**ENT, AUD, RSP and ORB**: in these cortical regions, (Herculano-Houzel, Watson et al. 2013) provides the bias-free neuron numbers generated from IF. The neuron density is obtained from dividing the neuron number by the CCFv3 regional volume. The subjects are four male C57/B6 mice aged 6 weeks.

RSP: 314,761/5.5147 ~ 57076 /mm^3

ENT: 400019/5.9712 ~ 67000/mm^3

AUD: 377362/3.03 ~ 124542/mm^3

ORB: 107179/3.09 ~ 34685 /mm^3

**CA1, CA3:** In (Fabricius, Wortwein et al. 2008), subjects are C57 2month 4 male and 8 female. Figure 8 provides the neuron number. We divide the number with CCFv3 volume.

In (Hlatky, Lui et al. 2003), the wild type C57/B6 subjects (male, age unspecified) provide the neuron density.

**CA1:** (Liu, Yu et al. 2006) SAMR1 (male, control, resistant to early senescence of age-related disease) mice in this study yield CA1 neuron number 206,000 ± 11,000 neurons in 4-month-old and 190,000 ± 10,000 in 8-month-old.

Density = 206000/CCF volume 5.38 = 38290 /mm^3 (4-month old)

BLA: (Mozhui, Hamre et al. 2007) we take the neuron number from C57/B6 subjects. The average age is 96 days and the range is from 30 to 500 days. The sex ration is 96 (F) : 103 (M).

**Image quality as a factor of segmentation failure**

The performance of our method is highly dependent on image quality. In cases where neurons are densely stacked or the noise is high, researchers may struggle to segment the neurons by eye. Unfortunately, some of the data used in our study suffers from these issues. To address this problem, our workflow includes a compromised approximation that estimates the blob's volume over the neuron volume when segmentation is challenging. This approximation is based on the median neuron volume from the training data. Figure S2 presents examples of cases where our workflow was unable to accurately segment neurons due to poor image quality and densely stacked neurons.

To avoid this issue, researchers can use higher quality data, such as higher resolution data from expansion microscopy or LSM acquired with a higher power objective. However, this solution may not fully address the problem, particularly in regions where neurons are densely packed.

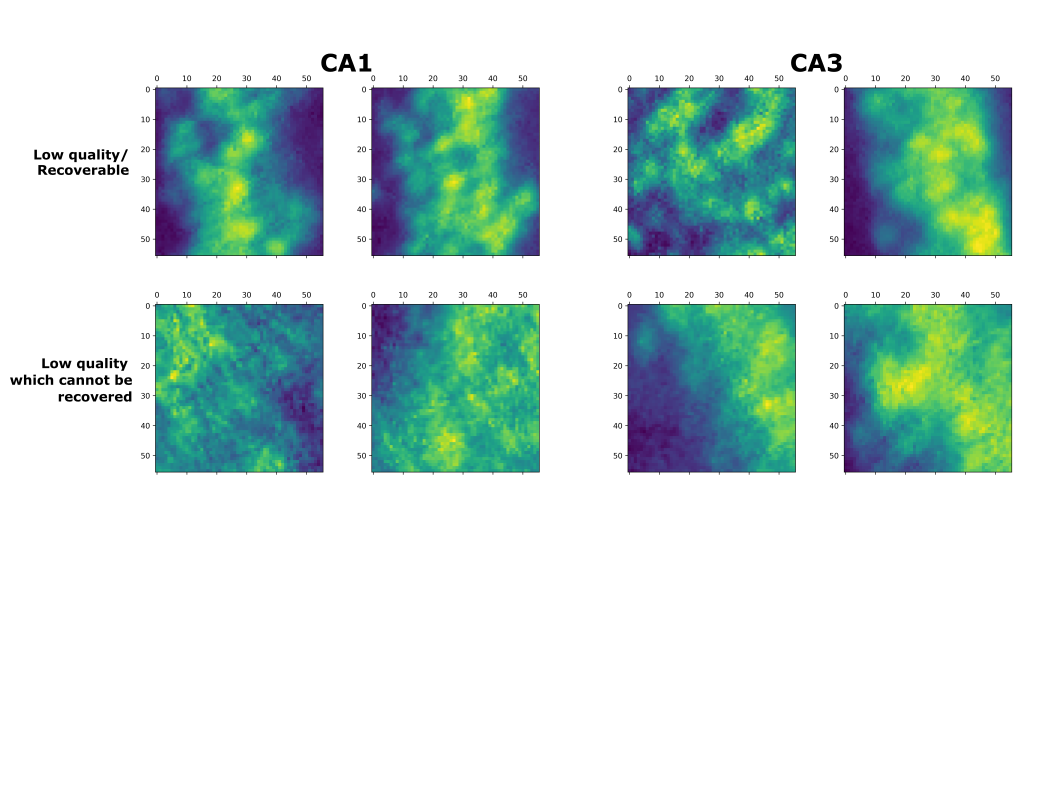

**Supplementary Figure 4.** provides examples of low-quality images due to densely stacked neurons or poor imaging conditions, taken from CA1 and CA3. The first row shows an image that can be counted using the approximation, while the second row shows an image with such poor quality that it cannot be recovered. The side scale shows the number of pixels, and each image measures 100 μm x 100 μm.

| Regions | Neuron density  (mean CV +/- SD) | Neuron number  (mean CV +/- SD) |
| --- | --- | --- |
| LGd | 0.11 | 0.13 |
| AUD | 0.08 | 0.08 |
| RSP | 0.07 | 0.06 |
| Orb | 0.09 | 0.07 |
| SUBR | 0.10 | 0.09 |
| VII | 0.33 | 0.35 |
| PSV | 0.24 | 0.44 |
| ENT | 0.15 | 0.13 |
| CA1 | 0.06 | 0.08 |
| CA3 | 0.09 | 0.07 |
| VISp | 0.15 | 0.13 |
| TH | 0.17 | 0.16 |
| BLA | 0.30 | 0.27 |

**Supplementary Table 2.** shows the coefficient of variation across animals.
